# Loss of *Regulator of telomere elongation helicase (Rtel1)* impairs alveolar epithelial cell progenitor function but does not exacerbate experimental fibrosis

**DOI:** 10.1101/2025.09.04.674361

**Authors:** Carla L. Calvi, Nichelle I. Winters, A. Scott McCall, Rafael J. Fernandez, Franck J. Kamga Gninzeko, Hao Ding, David S. Nichols, Vasiliy V. Polosukhin, Jennifer MS Sucre, Jason J. Gokey, Timothy S. Blackwell, Jonathan A. Kropski

## Abstract

Inherited loss-of-function mutations in the replication-associated helicase *Regulator of telomere elongation helicase* (*RTEL1*) are among the most frequent monogenic causes of familial pulmonary fibrosis (FPF), but the specific functional role of RTEL1 in lung development, homeostasis, and injury-repair has not been determined. Using single-cell RNA-sequencing to define the cellular diversity and molecular programs following intratracheal bleomycin injury in mice, we observed that *Rtel1* was most highly expressed in the lung epithelium, and its expression was reduced in PF epithelial cells compared to nonfibrotic controls. While global deletion of *Rtel1* leads to embryonic lethality, constitutive deletion of *Rtel1* in the lung epithelium (Rtel^Δepi^) led to normal lung structure and architecture. Rtel1^Δepi^ mice exhibited accelerated telomere attrition and impaired AEC progenitor potential *ex-vivo*. Despite decreased AEC progenitor function, no differences in lung fibrosis were observed in Rtel^Δepi^ mice compared to controls following single-dose or repetitive intratracheal bleomycin. Together, these data implicate RTEL1 as a regulator of AEC progenitor function but this level of progenitor dysfunction is not alone sufficient to cause or exacerbate experimental lung fibrosis.

## Introduction

Inherited loss-of-function mutations in genes related to telomere biology cause a spectrum of human disease, including dyskeratosis congenita, aplastic anemia, cryptogenic cirrhosis, and pulmonary fibrosis (PF) (1). Monoallelic inheritance of rare genetic variants in the telomere-related genes telomerase reverse transcriptase (*TERT*) (2, 3) and regulator of telomere elongation helicase (*RTEL1*) (4–6) are the most frequent monogenic causes of pulmonary fibrosis (7, 8), and available evidence suggest genetic risk for short telomeres contributes to IPF risk (9), but the mechanism by which telomere/telomerase dysfunction leads to progressive pulmonary fibrosis remains unclear.

Prior studies using telomerase reverse transcriptase (*Tert*) and telomerase RNA component (*Terc*) deficient mouse models have demonstrated that global loss of telomerase activity is not sufficient to cause spontaneous lung fibrosis in mice, while the impact of telomerase deficiency in experimental models of lung fibrosis has been inconsistent (10–12). Available evidence suggests that the impacts of telomerase mutations are mediated primarily through their effects on telomere length (13). However, the lung alveolar epithelium is a slow-turnover tissue (14, 15), thus it is uncertain as to why and how severe/critical telomere shortening occurs *in-vivo (16–18)* and the direct impact on cellular function have not been ascertained. Prior studies using genetic approaches to conditionally delete shelterin components (*Telomere repeat binding factor 1* and *2, Trf1* and *Trf2*, respectively) in alveolar epithelial cells indicate that widespread activation of a telomere DNA-damage response (DDR) can lead to acute lung inflammation and/or fibrosis (12, 19, 20), similar to other models of acute alveolar injury (21–26). However, the mechanism by which more indolent telomere attrition drives lung fibrosis has not yet been determined.

At the ends of each chromosome, telomere repeat sequences form a loop structure wherein the terminal ends of the 5’ and 3’ strands insert into the complementary telomere repea sequences, suppressing the sensing of the ends of each chromosome as double-stranded DNA-breaks (27). RTEL1 is a replication-associated iron-sulfur-cluster helicase responsible for unwinding the telomere D-loop to allow the telomerase holoenzyme access to telomere DNA sequences during cell division(28). Without RTEL1, during cell division the SLX4 resolvasome cleaves the T-loop, leading to rapid telomere attrition(28). In addition to its specific telomere-related function, RTEL1 has also been implicated as a regulator of replication stress (29, 30), genome stability (31–33), and prevention of transcription-replication collisions (34). These functions appear critical for normal development, as global deletion of Rtel1 leads to embryonic lethality in mice (35), and humans with biallelic loss-of-function mutations in *RTEL1* present early in life with severe Hoyeraal-Hreidarson syndrome (36–40).

We hypothesized that loss of Rtel1 specifically in the lung epithelium would lead to impaired alveolar epithelial regeneration following injury and promote experimental lung fibrosis. To our surprise, while epithelial deletion of *Rtel1* led to progressive telomere shortening in the lung epithelium and impaired AEC progenitor function in *ex-vivo* organoid models, this AEC dysfunction neither induced spontaneous lung fibrosis nor augmented fibrosis in multiple *in-vivo* models. These results imply that mechanisms beyond moderately impaired alveolar epithelial cell progenitor function are essential in linking genetic variation related to telomere biology to the development of pulmonary fibrosis.

## Materials and Methods

### scRNA-seq reanalysis

We queried previously published scRNA-seq data from 49 control and 65 PF lungs (GSE227136)(41) for expression of *RTEL1*. Data visualization was performed using Scanpy(42) v1.7.2. Subject-level pseudobulk differential expression at lineage-level was performed using deSEQ2 as previously reported (43). We also reanalyzed scRNA-seq data from unchallenged and bleomycin-treated C57Bl6 mice previously reported by our group (GSE243135)(44).

### Mice

All studies were approved by the IACUC at Vanderbilt University Medical Center. Sonic hedgehog (Shh)-cre mice(45), Sftpc-CreERT2(46), and UBC-CreERT2(47) mice were obtained from Jackson Laboratories (#005622, #028054, #007001), and crossed with *Rtel* floxed mice(35) (Rtel^f/f^) to generate Rtel^Δepi^, Rtel^ΔAT2^ and Rtel^iGKO^ mice. Mixed background Rtel^f/f^ mice were back-crossed onto a C57Bl/6 background for >7 generations prior to crossing with Cre lines. Cre-negative littermate animals were used as controls for all studies. All studies were approved by the Institutional Animal Care and Use Committee at Vanderbilt.

### Tamoxifen treatmen

Rtel^iGKO^, Rtel^ΔAT2^ mice and cre-negative littermate controls were administered tamoxifen citrate 400 mg/g diet (Envigo) x5 days on, 2 days off x 3 cycles followed by at least a 14 day washout prior to bleomycin injury.

### Histology

Experimental mice were euthanized by CO_2_ administration, the chest was exposed and descending aorta severed. Following cardiac puncture lungs were perfused with 10 ml cold PBS and inflated at 24 cm H_2_O with 10% neutral buffered formalin. Inflated lungs were placed in 50 ml conical tubes containing 25 ml buffered formalin and fixed for at least 24 hours prior to paraffin embedding as previously described (44, 48, 49).

### Telomere measurements

Telomere length quantification was performed by in situ hybridization utilizing the Telomere PNA FISH Kit/Cy3 (Dako K5326) following manufacturer’s instructions. Briefly, slides containing 5 µm sections of formalin fixed paraffin embedded lung tissue were rehydrated, and energy retrieved for 30 minutes at 100^0^C with Citrate buffer (Millipore 21545). Slides were allowed to cool to room temperature, washed thrice with diH_2_O followed by TBS, dehydrated in an Ethanol series and allowed to dry. Probe was applied to tissue and hybridized at 80^0^C, followed by 30 minutes at room temperature. Slides were rinsed and washed at 65^0^C for 5 minutes. After in-situ hybridization with PNA FISH, slides were rinsed twice with diH2O for 10 minutes followed by PBS,and blocked for 10 minutes with PowerBlock universal blocking reagent (Biogenex HK085-5K). An antibody mixture containing proSpC (Abcam 90716), HopX AF647(Santa Cruz SC398703), and SCGB1A1 AF488 (Santa Cruz 365992) diluted in 1%BSA/PBS was applied to hybridized tissue and incubated overnight at 4^0^C in an opaque humidified chamber. The following morning slides were washed three times with PBS 0.1% Triton and a donkey anti-rabbit dylight AF755 diluted in 0.1% Triton/PBS was applied to sections. Following a 30 minute incubation slides were again washed and nuclear detection achieved by a 10 minute incubation with Dapi dilactate. Slides were rinsed with PBS followed by diH2O and mounted with ProLong Gold (Thermofisher). After a cure period of 48-72 hours slides were sealed with nail polish and imaged in a Keyence BZ-X1710.

### Lung Morphometry

Lung morphometry was quantified using automated image analysis (AlveolEye, https://github.com/SucreLab/AlveolEye) as previously described (50).

### Bleomycin fibrosis models

Biweekly intratracheal bleomycin (McKesson 3224177; APP Pharmaceuticals 63323013720) was dissolved in sterile PBS and was administered by intratracheal instillation (single dose: 0.08 IU/100ul/mouse x1; repetitive: 0.04 IU/100ul/mouse x 4 to 8 doses) as previously described (24, 43, 44, 48, 49).

### Collagen quantification

At time of euthanasia the right lower lobe of experimental mice was excised, rapidly frozen in liquid nitrogen and stored at -80^0^C. Lung collagen was quantified with Sircol^TM^ kits (Biocolor, UK) following manufacturer’s instructions. Both soluble (S1000) and insoluble (S2000) fractions were measured. Briefly, the lobe was homogenized on ice with a 1.0ml solution of 0.1mg/ml Pepsin (Sigma Aldrich P7000) in 0.5M Acetic acid. Homogenates were incubated overnight at 4^0^C with gentle rocking. A 100 μl aqueous fraction of the homogenate was used to perform soluble collagen quantification, while the precipitated “residue” was used for insoluble collagen quantification. Collagen was quantified at 555 nm absorbance on a BioTek Synergy/LX multi-mode reader. For repetitive IT bleomycin studies, data are presented as pooled from two independent experiments, and total collagen is presented as a percent of the average of PBS-treated animals from each experiment to normalize for between-experiment batch-effects.

### Organoid generation

Lung tissue of aged (22/23 month old) Rtel^f/f^ and Rtel^Δepi^ mice (3 in each group) was digested with an enzyme cocktail containing 10IU Dispase II/1ml PBS, and resulting cell suspension was DNAse treated and strained thru 100µm followed by 40µm filters. Live cells were counted against trypan blue, and Type II AECS FACCS sorted utilizing the following parameters: Dapi negative or Calcein Violet AM positive, CD45/CD31/ter119 negative, CD326 and lysotracker positive. A minimum of 5×10^6^ cells for each animal was stained and FACCS sorted on a BD FACSAria III at the VUMC Flow Cytometry Shared Resource.

Type II sorted cells were counted, centrifuged and resuspended together with cultured ATCC MLG2908 mouse fibroblasts at a ratio of 5,000 AECII cells:25,000 MLG2908 in pre-warmed 200ul mix of MTEC plus media and Basement Membrane Extract Pathclear (Cultrex), 50% each, and cultured in 0.4µm transwell insert (Falcon, PET) placed in a 24 well plate. After allowing BME polymerization, 500ul MTEC plus/Retinoic Acid/KGF media containing ROCK inhibitor (Millipore Y-27632) was added to the lower chamber for the first 48 hours. Media in the lower chamber was changed 3 times weekly and cells were co-cultured for 14 days total. At the end of the co-culture period media was removed and cells were fixed with 4% paraformaldehyde in PBS overnight at 4^0^C, followed by PBS and stored at 4^0^C until imaged (Keyence BZ-X1710) for CFU quantification. After quantification membranes containing organoids encased in BME were carefully cut out, placed in biopsy cassettes in 10% buffered formalin for 24hrs and embedded in paraffin for histological analysis.

### Immunofluorescence

Slides containing 5 µm sections of formalin fixed paraffin embedded lung tissue or organoids were rehydrated, energy retrieved for 15-30 minutes at 100^0^C with Citrate buffer (Millipore 21545). Slides were allowed to cool to RT, washed twice with diH_2_O, treated for 20 minutes with 3% hydrogen peroxide in methanol(lung tissue), washed with diH2O followed by a 10 minute incubation in PBS. Tissue was permeabilized with 0.1% Triton in PBS for 10 minutes and blocked for 10 minutes with PowerBlock universal blocking reagent (Biogenex HK085-5K). An antibody mixture containing pro-SpC (Abcam 90716), and Hopx AF647(Santa Cruz SC398703), diluted in 1%BSA/10% Donkey Serum in PBS was applied to organoids and incubated overnight at 4^0^C in an opaque humidified chamber. The following morning slides were washed three times with PBS 0.1% Triton. Donkey anti-rabbit AF555 diluted in 0.1% Triton/PBS was applied to sections and allowed to incubate for 30 minutes. Slides were washed and nuclear detection achieved by a 10 minute incubation with Dapi dilactate. Slides were rinsed with PBS followed by diH2O and mounted with ProLong Gold (Thermofisher). After a cure period of 48-72 hours slides were sealed with nail polish and imaged in a Keyence BZ-X1710. Analysis was performed using Halo (Indica Labs) software.

### Data and code availability

No new genomic data were generated in this manuscript. Code used for plot generation is available at github.com/KropskiLab/Rtel1_2024.

### Statistics

Data are presented as mean +/- SEM or median +/- IQR depending on normality of the data unless otherwise specified in the legend. Mann-Whitney-U and Kruskal-Wallis with Tukey post-test adjustment for multiple testing were used for between/across group comparisons. P<0.05 was considered significant. Statistical analysis was performed using GraphPad Prism v9.0.

## Results

### RTEL1 is expressed in the regenerating lung epithelium

To first determine the cell types in which *RTEL1* is predominantly expressed, we interrogated single-cell RNA-sequencing data from control and IPF lungs(41). *RTEL1* expression was generally expressed in most cell types but at very low levels in IPF and control lungs (**Figure 1A**). Performing subject-level pseudobulk differential expression analysis, RTEL1 express was nominally lower in Epithelial and Proliferating Epithelial cells (but not immune, endothelial or mesenchymal cells) from IPF lungs compared to controls (**Figure 1B**), although these differences did not reach multiple testing-adjusted significance in global differential expression analyses. Next, we reanalyzed plate-based scRNA-sequencing data of C57Bl6 mice following IT bleomycin or PBS(44) (**Figure 1C)**. Similar to what we observed in the human lung, *Rtel1* was expressed at low levels in most cell types; as anticipated, expression was highest in proliferating cells, and there was low-level expression in the lung epithelium (**Figure 1C**). Given the well-established links of the lung epithelium to PF susceptibility(51) and the postulated roles of telomere-related genes epithelial progenitor function/senescence, we focused our studies on the role of *Rtel1* in epithelial cells.

**Figure 1.**
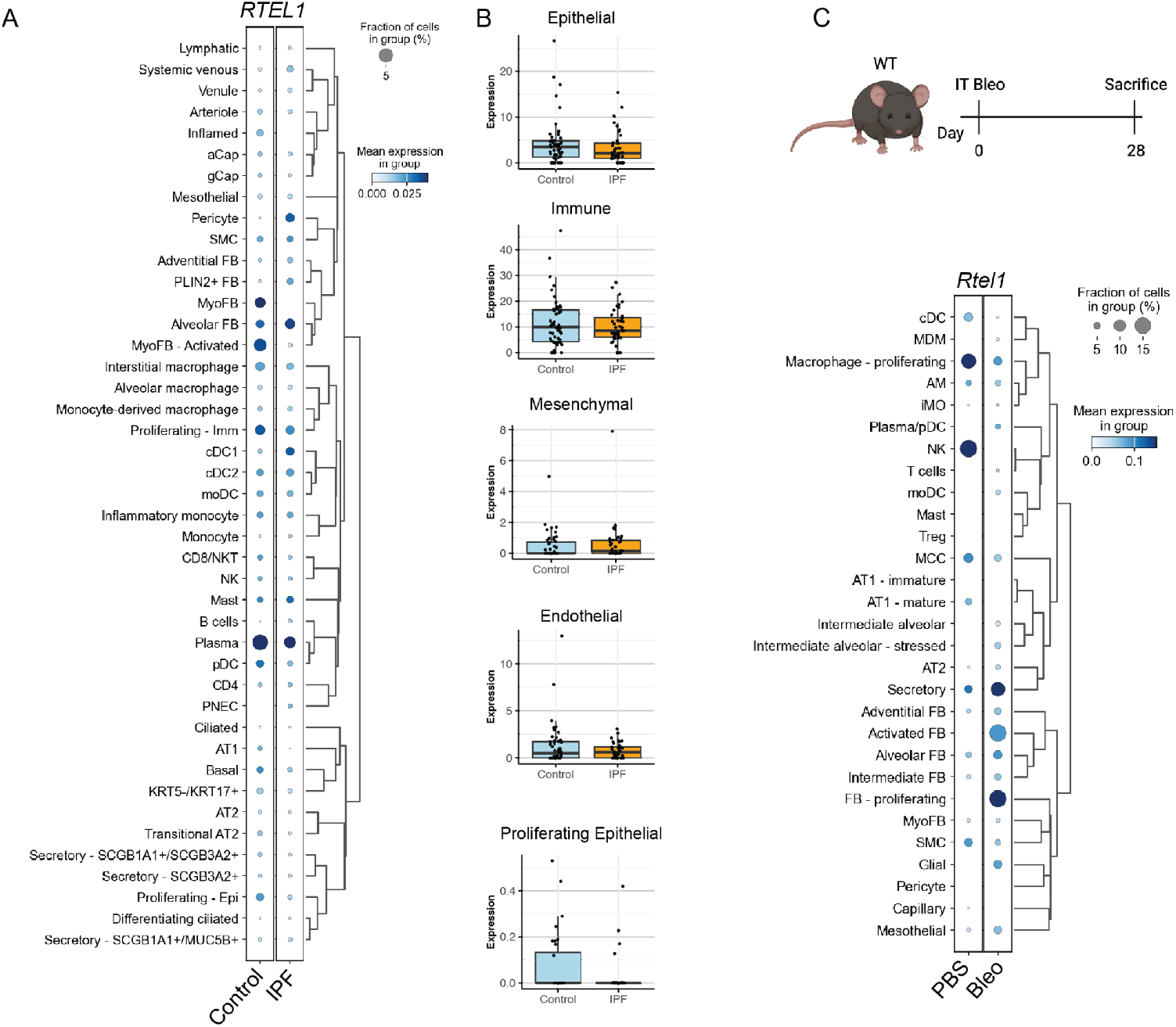
*RTEL1* expression in the human and mouse lungs. A) Dotplot depicting expression of *RTEL1* in control and IPF lung samples (from GSE227136). B) Subject level pseudobulk differential expression analysis performed at the lineage level from GSE227136. Assessment of differential expression did not reach statistical significance after transcriptome-wide multiple testing adjustment. C) Dotplot of *Rtel1* expression in C57Bl6 mice at day 28 after bleomycin injury (from GSE243135).

### Epithelial Rtel1 deletion accelerates telomere attrition without altering normal lung development

To investigate the role of Rtel1 in the lung epithelium, we generated pan-lung epithelial Rtel1 deficient mice (*Shh-Cre; Rtel1*^*f/f*^, Rtel1^Δepi^ hereafter). The Shh-Cre is activated in lung bud progenitors within 1 day of lung-specification (approximately embryonic day 9.5, E9.5), has high (>99%) efficiency, ensuring that entire lung epithelium is derived from recombined (Rtel1 deficient) cells(45, 52). In contrast to what was observed for global Rtel null mice (35), Rtel1^Δepi^ mice were viable, born in normal Mendelian ratios and exhibited norma-appearingl lung morphology (**Figure 2A-B**). We then sought to determine whether Rtel1 deletion would lead to lineage-specific telomere length attrition in adult mice (4 months of age). We performed multiplexed immunofluorescence/ISH co-staining for canonical distal-lung epithelial lineage markers Scgb1a1 (secretory cells), pro-Spc (AT2 cells) and Hopx (AT1 cells) together with a telomere PNA probe (**Figure 2C**) and quantified lineage-specific telomere length by automated image analysis (see Methods). We observed significantly shorter telomeres in AT2 cells from Rtel1^Δepi^ mice compared to controls, however telomere length was similar between groups in secretory and AT1 cells (**Figure 2D-F**). Regardless of genotype, telomere length was longest in secretory cells and shortest in AT1 (**Figure 2D-F**). These data imply that deletion of Rtel1 leads to accelerated telomere attrition in AT2 cells.

**Figure 2.**
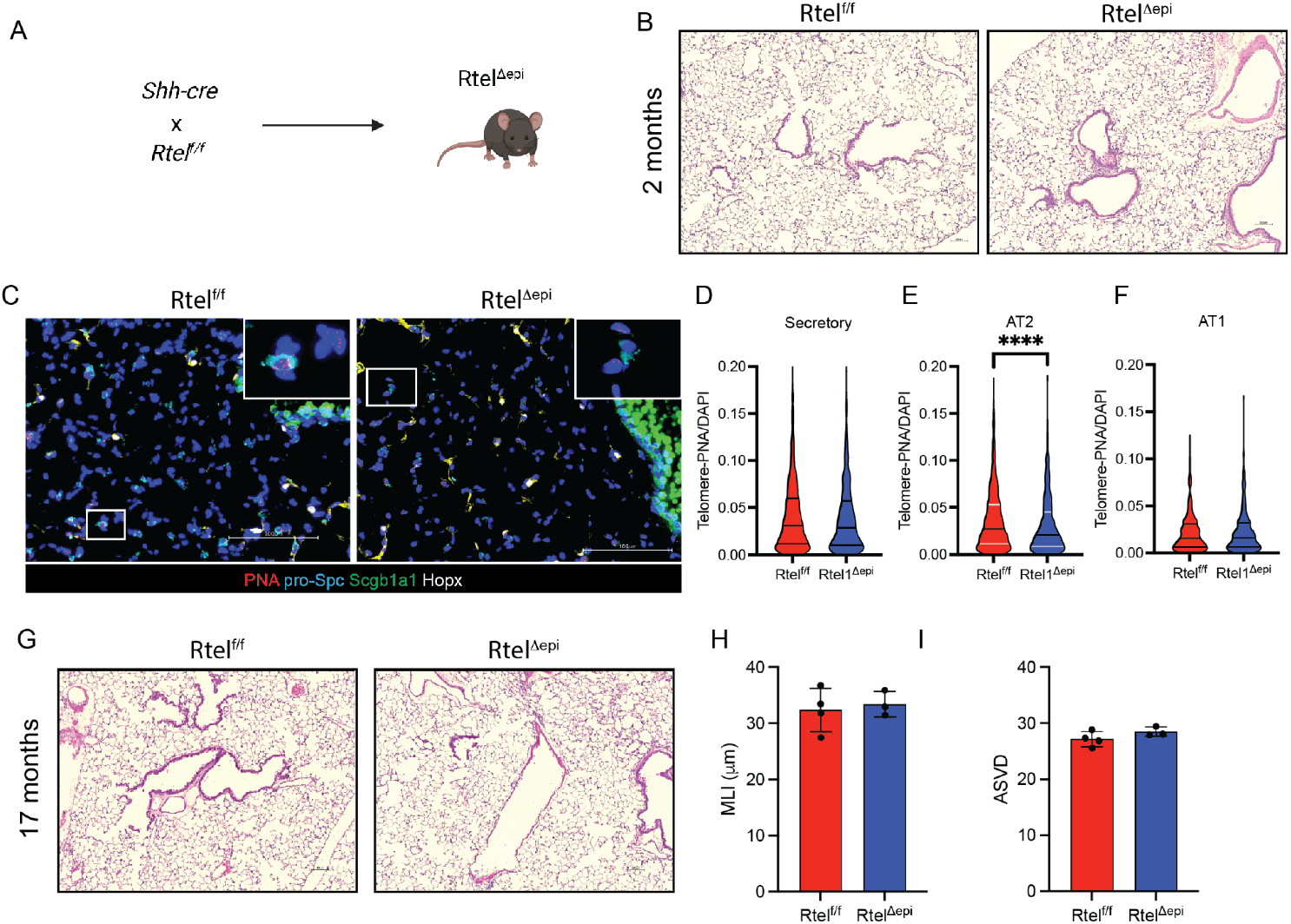
Deletion of Rtel1 in the lung epithelium leads to telomere shortening in AT2 cells without altering lung morphometry. A) Schematic of Rtel1^Δepi^ mouse generation. B) Representative H&E images from 2 month old Rtel1^Δepi^ and littermate control mice. C) Representative IF/ISH images co-staining for telomere-PNA and epithelial lineage markers in Rtel1^Δepi^ and Cre-negative littermates at 4 months of age. Insets are enlargements of highlighted white boxes indicating a representative AT2 cell. Quantification of epithelial cell telomere length in D) Secretory, E) AT2 and F) AT1 cells. **** P<0.0001 by Kruskall-Wallis. G) Representative H&E images of unchallenged Rtel1^Δepi^ and Cre-negative littermates 17 months of age. Quantification of (H) mean-linear intercept (MLI) and (I) airspace density volume (ASVD) by automated image analysis. Scale bars = 100 μm.

We next asked whether this accelerated telomere attrition would lead to aging-related changes in lung structure. Rtel1^Δepi^ and cre-negative littermate controls were aged to ∼17 months (**Figure 2G**) and lung morphometry was assessed by automated image analysis. Mean linear intercept (**Figure 2H**) and airspace volume density (ASVD, **Figure 2I**) were similar between groups. These results suggested that while loss of *Rtel1* leads to shortened AT2 cell telomeres, this does not lead to spontaneous fibrosis or emphysema.

### Rtel1 promotes AEC progenitor function and differentiation

We then sought to determine whether the accelerated telomere shortening in AT2 cells would alter AT2 progenitor function. To test this, we sorted Cd45-/Cd31-/Epcam^+^/Lysotracker^+^ epithelial cells from Rtel1^Δepi^ and age-matched cre-negative littermate controls by FACS and established coculture organoids with MLG2908 fibroblasts (**Figure 3A**). We found that AECs isolated from Rtel1^Δepi^ mice generated fewer organoids compared to controls (**Figure 3B-C**). Immunofluorescence analysis demonstrated fewer Hopx^+^ cells per organoid generated from Rtel1^Δepi^ mice compared to controls (**Figure 3D-E**) (indicating reduced AT2 -> AT1 differentiation). Together, these data suggested that deletion of Rtel1 impairs proliferation and differentiation of AT2 cells.

**Figure 3.**
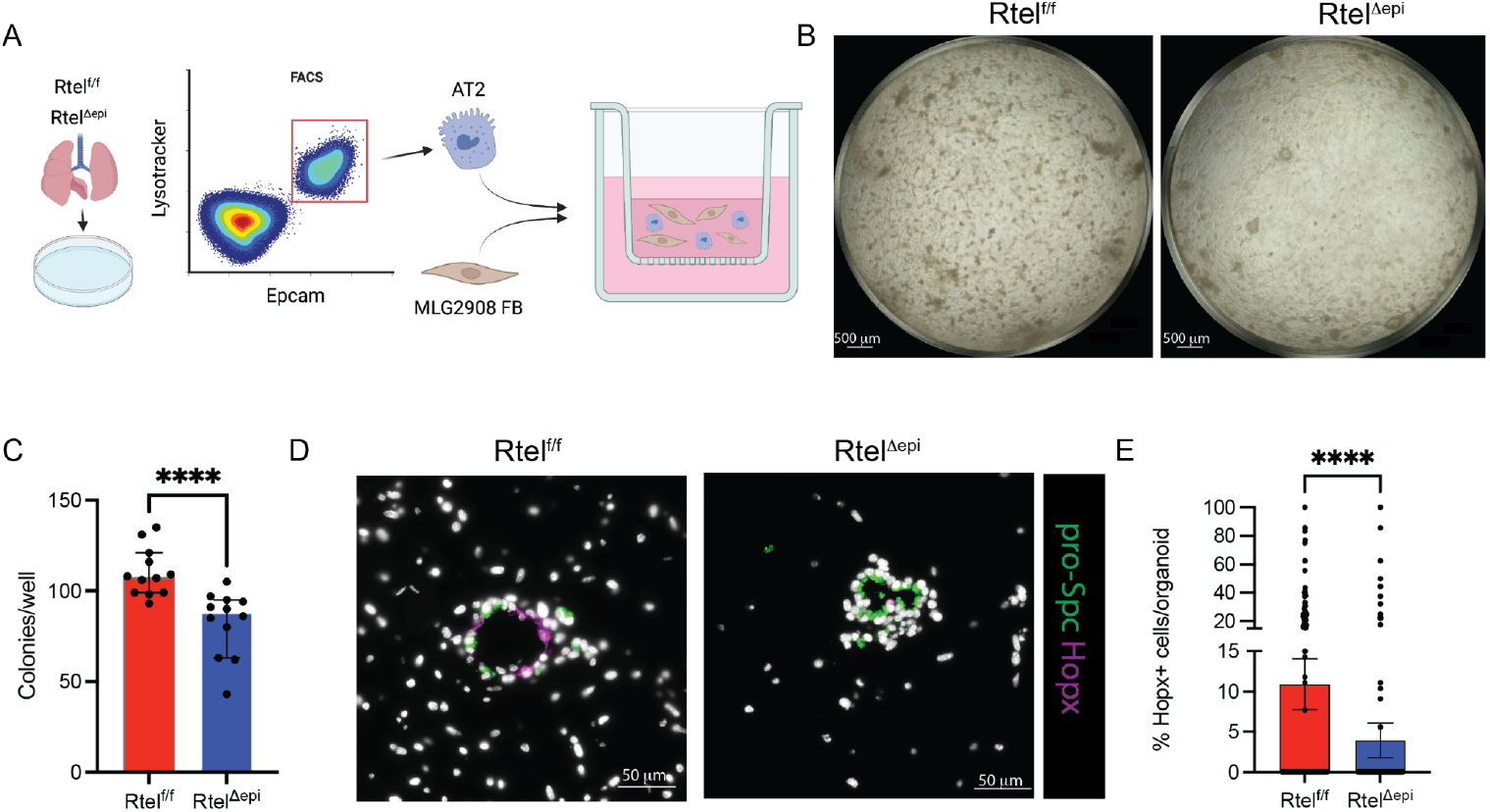
Epithelial Rtel1 deletion reduces AT2 progenitor function *in-vitro*. A) Schematic summarizing alveolar organoid generation. Single cell suspections were generated from lung tissue of unchallenged Rtel1^Δepi^ and Cre-negative littermates, and AT2 cells were isolated by FACS (Cd45^-^/Cd31^-^/Cd326^+^/Lysotracker^+^), and cocultured with MLG2098 fibroblasts (5K AT2: 25K fibroblasts) in 50% matrigel in MTEC^+^ media for 14 days. B) Whole-well images of organoids after fixation on day 14. C) Quantification of colonies in Rtel1^Δepi^ and Cre-negative littermates. D) Immunofluorescence staining of alveolar organoids for AT2 (pro-Spc) and AT1 (Hopx) markers. E) Quantification of Hopx^+^ cells per organoid in Rtel1^Δepi^ and Cre-negative littermates. **** P<0.0001 by Kruskal-Wallis.

### Deletion of Rtel1 in the lung epithelium does not alter fibrotic susceptibility

With evidence that loss of Rtel1 accelerates age-related telomere attrition and this is accompanied by reduced progenitor function in AECs, we then sought to determine whether loss of Rtel1 would exacerbate experimental lung injury and fibrosis. Our group and others have previously reported that the repetitive intratracheal bleomycin model leads to more persistent lung fibrosis in mice compared to single-dose bleomycin, and we hypothesized that repeated injury would be most suited to test the telomere-dependent replicative capacity for AECs on fibrotic remodeling. We aged Rtel1^Δepi^ and cre-negative littermate control mice to ∼8 months, then challenged them with biweekly IT bleomycin (0.04 IU every 2 weeks x 6 cycles) (**Figure 4A**) and sacrificed them for assessment of lung fibrosis 2 weeks after the final bleomycin dose (∼1 year of age). To our surprise, there were no differences in histologic fibrosis or lung collagen content comparing Cre-negative littermates and Rtel1^Δepi^ mice (**Figure 4B-C**). Similar results were observed in the single-dose IT bleomycin model using either the Rtel1^Δepi^ or AT2-specific Rtel deletion using the Sftpc-CreER (Rtel^ΔAT2^) model (**Supplemental Figure 1**).

**Figure 4.**
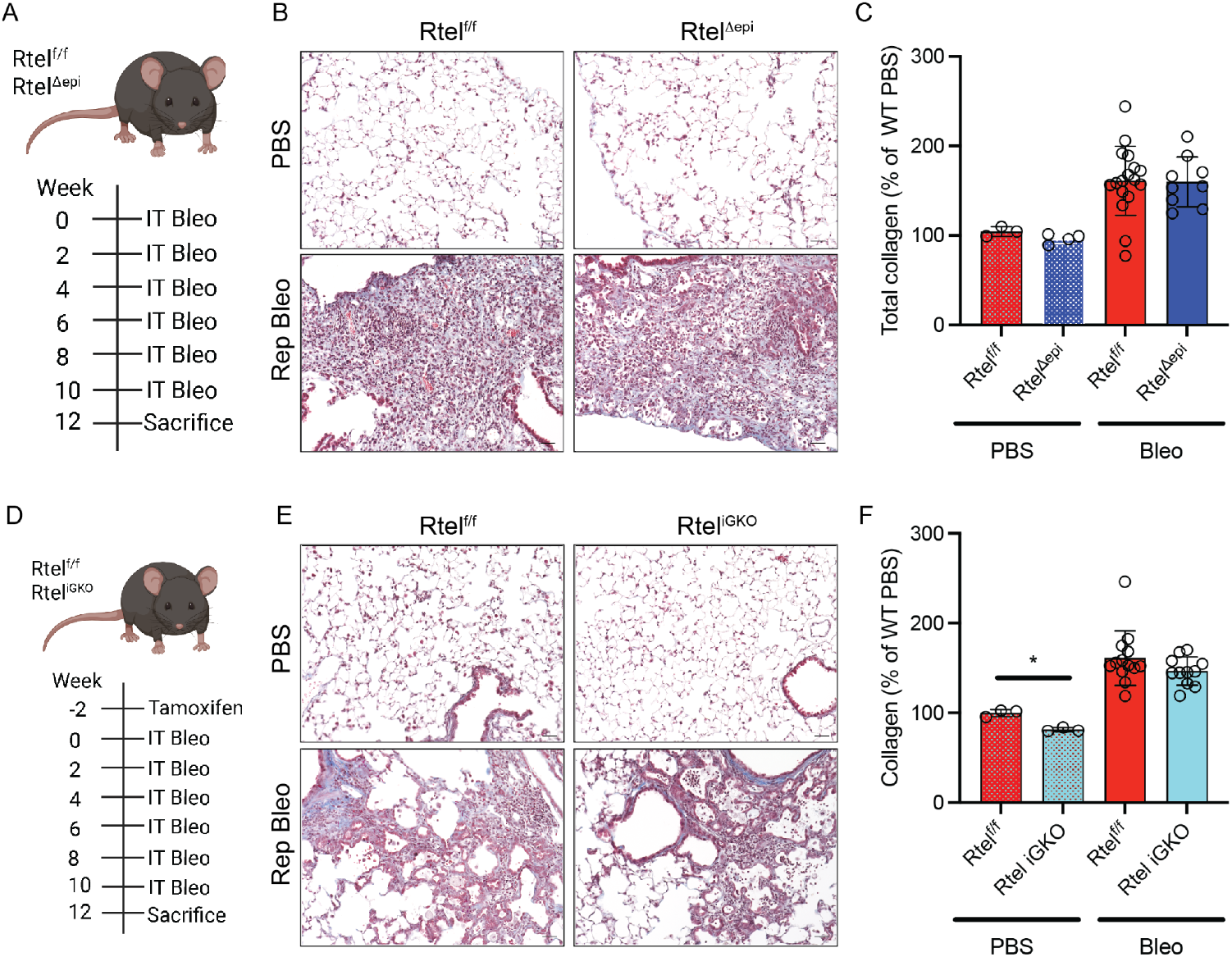
Experimental lung fibrosis is not altered by *Rtel1* deletion. A) Schematic of repetitive IT bleomycin model in Rtel1^Δepi^ and control mice. B) Representative Masson Trichrome images from PBS control and repetitive bleomycin (Rep Bleo) treated mice. C) Quantification of total lung collagen as measured by Sircol assay. Data represent pooled results from two independent experiments and are presented normalized to WT PBS controls from the given experiment. D) Schematic of repetitive IT bleomycin model in Rtel1^iGKO^ and control mice. E) Representative Masson Trichrome images from PBS control and repetitive bleomycin (Rep Bleo) treated mice. F) Quantification of total lung collagen as measured by Sircol assay. Data represent pooled results from two independent experiments and are presented normalized to WT PBS controls from the given experiment. Scale bars = 100 μm.

Since Rtel1 can be expressed not only in epithelial cells, we then wondered whether the Rtel1 primarily mediated its effects on fibrosis through other cell types. To test this, we generated Rtel1 global-inducible knockout mice (Rtel^iGKO^) using the ubiquitin-c (UBC) creER (**Figure 4D**). Compared to tamoxifen-treated cre-negative littermates, Rtel^iGKO^ mice again had similar histologic fibrosis and lung collagen content (**Figure 4E-F**). Together, these data suggest that telomere-length independent roles of Rtel1 do not contribute substantially to fibrotic susceptibility or lung fibrogenesis.

## Discussion

Genetic evidence strongly supports the concept that telomere biology is central to risk for pulmonary fibrosis, and loss-of-function variants in *RTEL1* appear to be the second most frequent monogenic cause of Familial Pulmonary Fibrosis(7, 8). The prevailing conceptual hypothesis linking telomere dysfunction to pulmonary fibrosis risk suggests that telomere shortening leads to replicative senescence in alveolar epithelial cells, resulting in failure of injury-repair (53), yet prior studies using genetic mouse models of telomerase deficiency (either *Tert* or *Terc* null mice) have yielded inconsistent results(10–12). Through its DNA-helicase function, loss of RTEL1 function can lead to rapid telomere shortening and activation of a telomere DNA-damage response(28), which we hypothesized may overcome some of the prior limitations studying lung telomere biology in mice. In this study, we found that epithelial deletion of *Rtel1* leads to telomere attrition in AT2 cells that is associated with reduced AT2 progenitor capacity, but this alveolar epithelial progenitor defect was not sufficient to cause or exacerbate lung fibrosis. Further, global deletion of Rtel1 in adult mice did not alter fibrotic susceptibility, implying that it is less likely that a telomere-length independent mechanism is a substantial contributor to the role of *Rtel1* in fibrogenesis.

These results add to a complex milieu of transgenic mouse modeling approaches of telomere dysfunction in the context of pulmonary fibrosis. Previous studies of *Tert* and *Terc* null mice have yielded contradictory results (10–12); our group previously found that *Tert* or *Terc* deficiency neither protected against nor exacerbated bleomycin-induced fibrosis, even when challenged with repeated injury, aged to 18+ months, and bred through multiple generations reaching reproductive failure(11). An overarching question in these studies is whether lung (AEC) telomere length would shorten sufficiently to trigger a telomere-DDR; this has generally not been observed spontaneously in these models using C57Bl6 mice which have very long telomeres at baseline. In contrast, models of shelterin component (Trf1 or Trf2) deletion (triggering a DNA-damage response) in AT2 cells consistently lead to lung injury, inflammation, and/or fibrosis (12, 19, 20).

We speculated that Rtel1 deficiency would accelerate telomere attrition and could have the potential for AECs to reach critically short thresholds more rapidly without life-limited systemic/reproductive impacts. Consistent with this hypothesis, we found that lung epithelial Rtel1 deletion does lead to telomere shortening in AT2 cells, and that this is associated with a proliferative defect. The observation that differences in epithelial cell telomere length manifest first in AT2 cells (which are slow turnover) rather than airway secretory cells (which have more rapid turnover) is unexpected. Further, the observation of similar (and short) telomere length in AT1 regardless of Rtel1 sufficiency suggests that AEC maturation is associated with telomere attrition, and that this mechanism may be replication independent.

Despite this progenitor defect, whether modeled using a single acute injury or repeated injury, epithelial Rtel1 deletion did not alter fibrosis, a finding that contrasts with more severe models of AT2 progenitor dysfunction (25, 54). One potential explanation is that the degree of telomere shortening observed (and effects on ex-vivo progenitor potential), while statistically significant, was insufficient to translate to marked *in-vivo* effects on fibrosis. These results do not exclude the possibility that more severe telomere shortening could be required to more influence in-vivo epithelial repair and fibrosis. Whether relative preservation of telomere length is due to the slow turnover of the alveolar epithelium, activation of the alternative lengthening of telomeres (ALT) pathway, or other factors is not entirely clear. Taken together, these results imply that inhibiting AT2 progenitor function (moderately) is insufficient to cause or exacerbate lung fibrosis.

Recognizing that fibroblast senescence (55) and short telomeres in circulating immune cells (16) have also been linked to PF, we also considered the possibility that other cell types (such as myeloid cells or fibroblasts) might mediate the effects on Rtel1 on fibrogenesis. Using a global inducible deletion strategy (which overcomes the embryonic lethality observed in *Rtel1* null mice (35), but also does not impact telomere length substantially in slow-turnover tissues), we again found that Rtel1 deletion neither protected against nor exacerbated experimental lung fibrosis.

This study has a number of limitations, the most pertinent of which is that inbred C57Bl6 mice have long telomeres at baseline; emerging models with more human-relevant telomere lengths (56) may provide new opportunities to interrogate these mechanisms *in-vivo*. We focused our analyses around measurements of peak fibrosis; it is possible that Rtel1 deficiency could alter fibrosis resolution, although we consider this less likely given the findings we observed in multiple repeated injury models.

To summarize, epithelial *RTEL1* deletion leads to telomere shortening and a proliferative defect in AT2 cells, these changes were insufficient to cause or exacerbate lung fibrosis in both acute and repeated injury models, highlighting the complexity of telomere dysfunction’s contribution to fibrogenesis and the need for more human-relevant models to advance understanding.

## Author contributions

CLC performed experiments, analyzed data, wrote and revised manuscript. NIW, ASM, RJF, FJK, DSN performed experiments and analyzed data; HD provided mice; VVP, JMSS, JJG analyzed data and revised manuscript, TSB designed study, provided oversight and revised manuscript, JAK designed study, performed experiments, analyzed data, wrote and revised manuscript.

## Supplementary Figures

**Figure S1.**
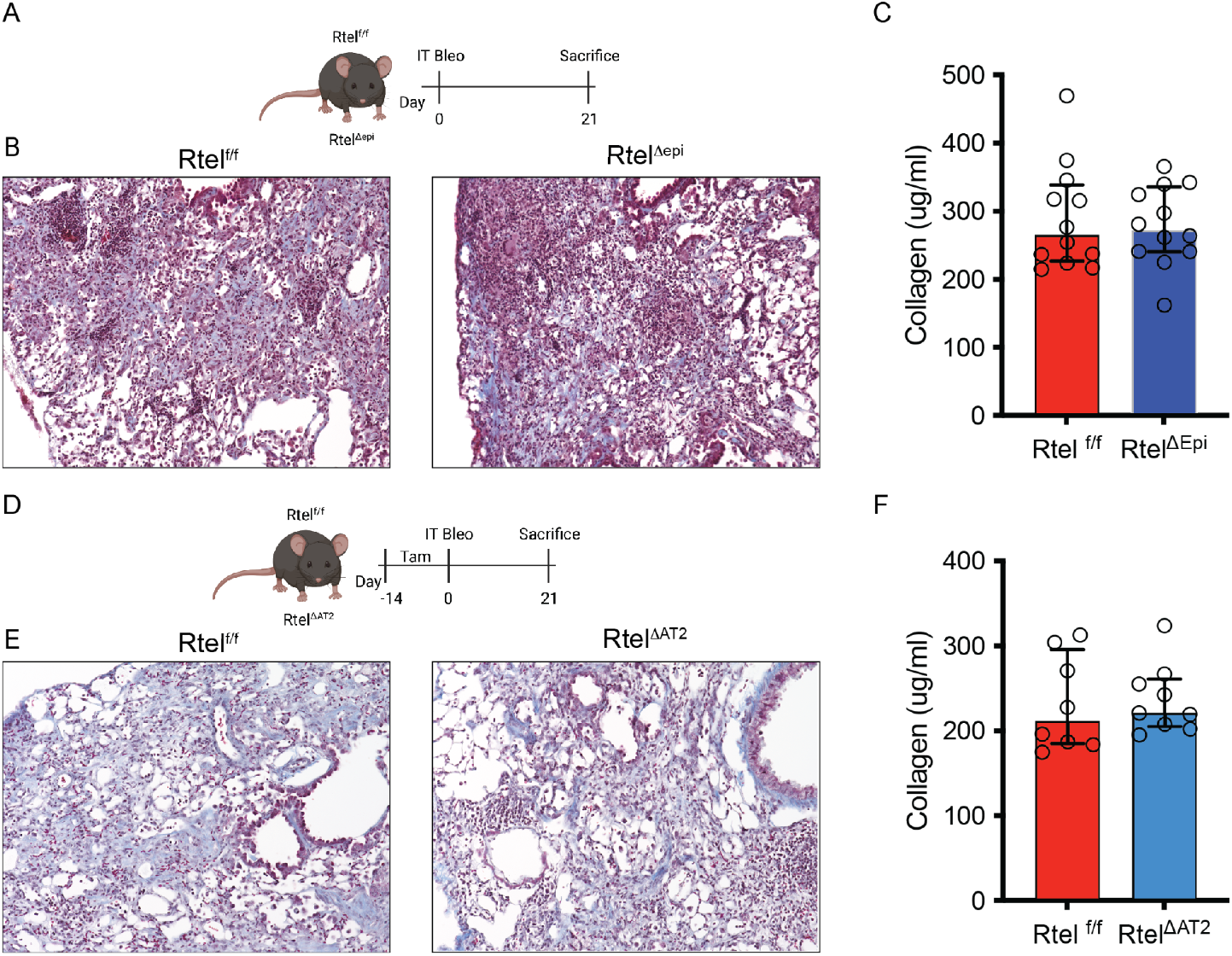
Pan lung epithelial and AT2-specific deletion of Rtel1 does not alter single-dose bleomycin-induced fibrosis. A) Schematic of single-dose IT bleomycin model in Rtel1^Δepi^ and control mice. B) Representative Masson Trichrome images and bleomycin treated mice. C) Quantification of total lung collagen as measured by Sircol assay. D) Schematic of single-dose IT bleomycin model in Rtel1^ΔAT2^ and control mice. E) Representative Masson Trichrome images from bleomycin treated mice. F) Quantification of total lung collagen as measured by Sircol assay.

